# Time-of-day influences on skill acquisition and consolidation after physical and mental practices

**DOI:** 10.1101/2021.11.02.466949

**Authors:** Charlène Truong, Pauline M. Hilt, Fatma Bouguila, Marco Bove, Florent Lebon, Charalambos Papaxanthis, Célia Ruffino

**Author notes:** **Corresponding Author:** TRUONG Charlène, Université de Bourgogne Franche-Comté, UFR STAPS, Campus Universitaire, BP 27877, F-21000 Dijon, France. Equal contribution (Charlène Truong, Pauline M. Hilt, Charalambos Papaxanthis).

## Abstract

Time-of-day influences both physical and mental performance. Its impact on motor learning is, however, not well established yet. Here, using a finger tapping-task, we investigated the time-of-day effect on skill acquisition (i.e., immediately after a physical or mental practice session) and consolidation (i.e., 24 hours later). Two groups (one physical and one mental) were trained in the morning (10 a.m.) and two others (one physical and one mental) in the afternoon (3 p.m.). We found an enhancement of motor skill following both types of practice, whatever the time of the day, with a better acquisition for the physical than the mental group. Interestingly, there was a better consolidation for both groups when the training session was scheduled in the afternoon. Overall, our results indicate that the time-of-day positively influences motor skill consolidation and thus must be considered to optimize training protocols in sport and clinical domains to potentiate motor (re)learning.

## Introduction

Motor learning is essential to acquire new and improve existing motor skills in a constantly changing environment. Exhaustive training is necessary to attain a rich motor repertoire on a long-term scale. Initially, the acquisition of motor skills is fast; a single or a few training sessions are sufficient to enhance accuracy and/or speed in a motor task ^1^. Although practice is undeniable for motor skill acquisition, its consolidation, namely the transformation of a new initially fragile motor memory into a robust and stable motor memory, can occur during rest periods without additional practice (off-line learning process). Indeed, several studies highlighted the fundamental role of sleep and the passage of time on motor skill consolidation ^2–4^.

Physical practice (PP) is not the only way to acquire or improve a motor skill. Mental practice (MP), which is the mental simulation of an action without any corresponding motor output, is a widespread form of training with proven efficacy in motor learning ^5,6^. MP improves muscle strength and flexibility ^7,8^, as well as speed and accuracy ^3,5,9^, representing thus a promising method for motor rehabilitation ^10^ and sports performance ^11^. The positive effects of MP on skill acquisition can be explained by the common properties it shares with PP ^12,13^. Motor learning through MP is associated with neural activations at several levels within the central nervous system, such as the parietal and prefrontal cortices, the supplementary motor area, the premotor and primary motor cortices, the basal ganglia, and the cerebellum. Recently, we have shown that during MP, the generation of a subliminal motor command triggers both cortical and subcortical circuits. This activation induces plastic modulations leading to important gains in motor performance ^14,15^. Moreover, after MP, the off-line learning process was also highlighted, leading to a more robust motor memory ^16^.

Current knowledge on motor learning leads to the assumption that the key for the formation of a rich motor repertoire can be found in the intelligent combination of periods with practice (physical or mental) and with rest (off-line learning). The elaboration of training programs, however, should also consider the time-of-day in which practice takes place. Indeed, several studies have suggested that physical and mental performances fluctuate through the day, on a circadian basis (∼24 h). For example, circadian variations have been observed for the maximal voluntary contraction ^17^, as well as for simple motor tasks, such as handwriting ^18^ and counter-flicking target performance ^19^. Likewise, more athletic movements also exhibit this circadian rhythmicity, such as in length of jump ^20^, the accuracy of badminton or tennis services ^21,22^, and the swimming speed ^23^. Generally, studies showed better performances in the afternoon than in the morning ^24,25^. Interestingly, Gueugneau et al. showed daily fluctuations in the timing of both physical and mental arm movements ^26–28^. Brain activation during physical and mental movement also shows strong circadian variations ^29^. Precisely, a contrast fMRI analysis revealed greater activity in the cerebellum, the left primary sensorimotor cortex, and the parietal lobe in the morning than in the afternoon during physical movements. The same analysis for the mental movement revealed increased activity in the left parietal lobe in the morning than in the afternoon. The reduction of cerebral activity in the afternoon could be related to the improved efficiency of the recruited neural circuits.

Although many studies have enriched the literature about the time-of-day influence on motor and mental performance, its impact on motor learning remains up to now unknown. The current study aims to evaluate the influence of the time-of-day on the acquisition and consolidation processes following PP and MP. According to daily fluctuations of physical and mental performances (morning vs. afternoon, Gueugneau et al. 2010), we scheduled two morning groups at 10 a.m. (G10_PP_ and G10_MP_) and two afternoon groups at 3 p.m. (G3_PP_ and G3_MP_), on two consecutive days. On day 1, we used a finger tapping task^2^ to measure the acquisition process (i.e., the improvement in skill performance immediately after PP or MP). On day 2, we measured the consolidation process on the same task (i.e., the improvement or stabilization in skill performance 24h after PP or MP). Following the existing literature, we hypothesized a greater gain after PP than MP ^3,5,30,31^. Due to the known variations of the physical and mental performances within a day, we hypothesized a better acquisition in the afternoon (whatever the mode of practice) than in the morning. In the absence of previous data on the consolidation process and time-of-day, we expected it to follow the same trend as acquisition, i.e., better consolidation of the motor skill after training in the afternoon than in the morning.

## Results

Forty-eight right-handed healthy adults were requested to tap on a computer keyboard an imposed sequence with their left hand (Fig. 1a). The participants were randomly assigned into four groups: two PP and two MP groups, trained in the morning (at 10 a.m., G10_PP_ and G10_MP_) and in the afternoon (at 3 p.m., G3_PP_ and G3_MP_) on day 1 (Fig. 1b). To evaluate the improvement in skill performance (i.e., the acquisition process), all groups physically accomplished the first two trials (1 and 2, pre-test, T1) and the last two trials (47 and 48, post-test, T2). The remaining trials (3-46, n = 44) constituted the training trials for the physical (G10_PP_ and G3_PP_) or the mental (G10_MP_ and G3_MP_) groups. To measure the consolidation process, all participants physically accomplished two trials 24 hours later (on day 2, T3). We recorded the accuracy and speed of the sequence execution and defined the motor skill as the combination of both (see Fig. 1a).

**Figure 1.**
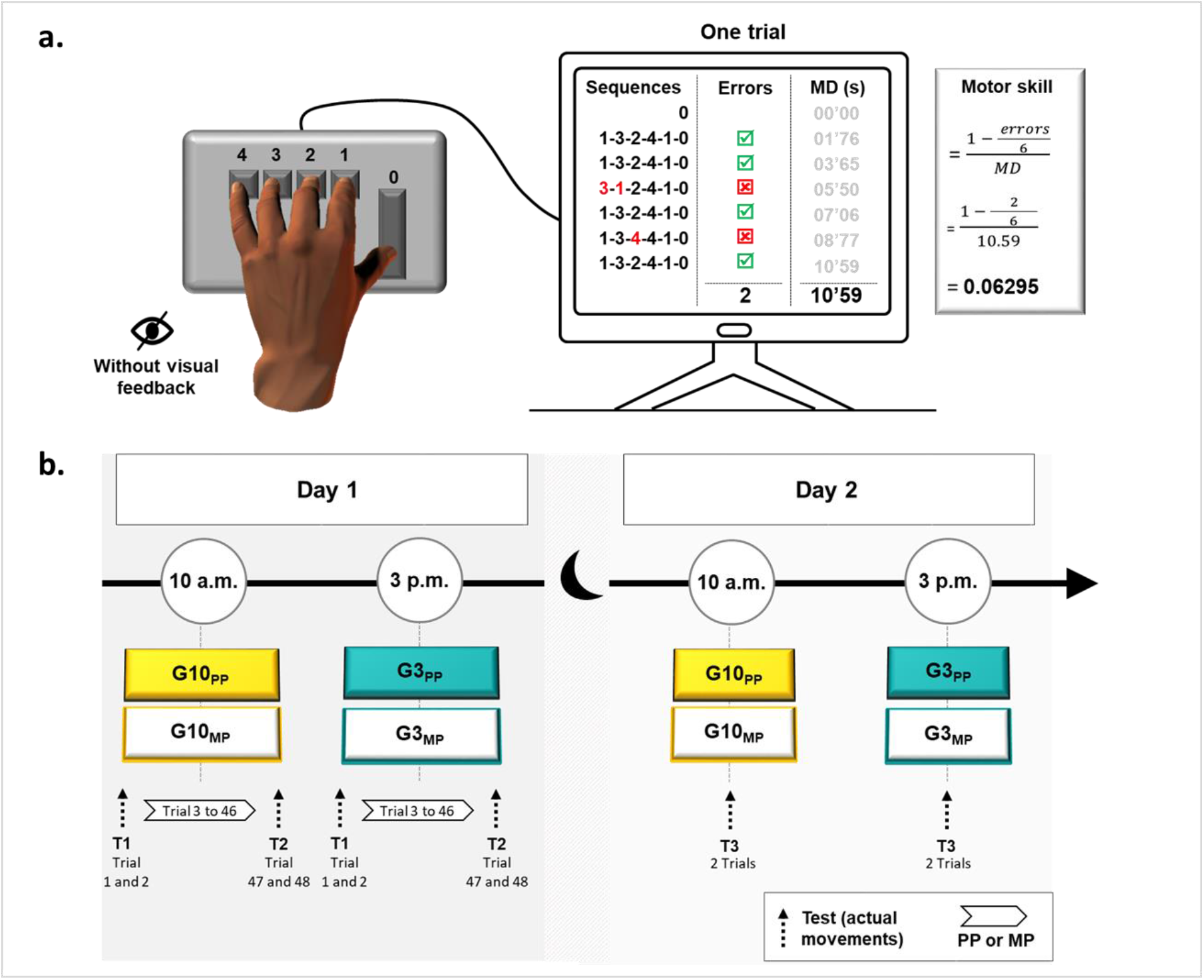
Illustration of experimental device and procedure. **(a)** The computerized version of the sequential finger-tapping task. Each key was affected to a specific finger of participants’ left hand: 0 (thumb), 1 (index), 2 (middle), 3 (ring), and 4 (little). Participants were requested to tap the following sequence as accurately and as fast as possible: 1-3-2-4-1-0. Six consecutive sequences composed one trial. Accuracy was defined as the number of false sequences (Errors) throughout one trial. Movement duration (MD) was defined as the time interval between the start of the trial (the first pressure on the key ‘0’) and the end of the trial (the last pressure on the key ‘0’, at the end of the 6th sequence). Motor skill is a composite ratio of duration and accuracy. **(b)** Experimental procedure. The participants were divided into four groups: G10PP physically trained at 10 a.m., G10MP mentally trained at 10 a.m., G3PP physically trained at 3 p.m., and G3MP mentally trained at 3 p.m. The protocol was scheduled on two consecutive days. The Day 1, participants trained on 48 trials: the two first trials and the last two trials were physically performed and composed T1 and T2, respectively. The remaining 44 trials constituted physical or mental practice. The Day 2, participants physically performed two trials 24 hours later (T3).

### Motor skill

Figure 2 shows the average values (+SE) of skill performance for the four groups (G10_PP_, G10_MP_, G3_PP_, and G3_MP_) and the three sessions (T1, T2, and T3). ANOVA revealed a significant interaction effect (F_6,88_ = 3.42, p < 0.01, η^2^ = 0.19). The post-hoc analysis showed similar initial skill levels (in all, p > 0.89) with comparable movement accuracy and duration (see, respectively, error rate and movement duration in Table 1).

**Table 1.**
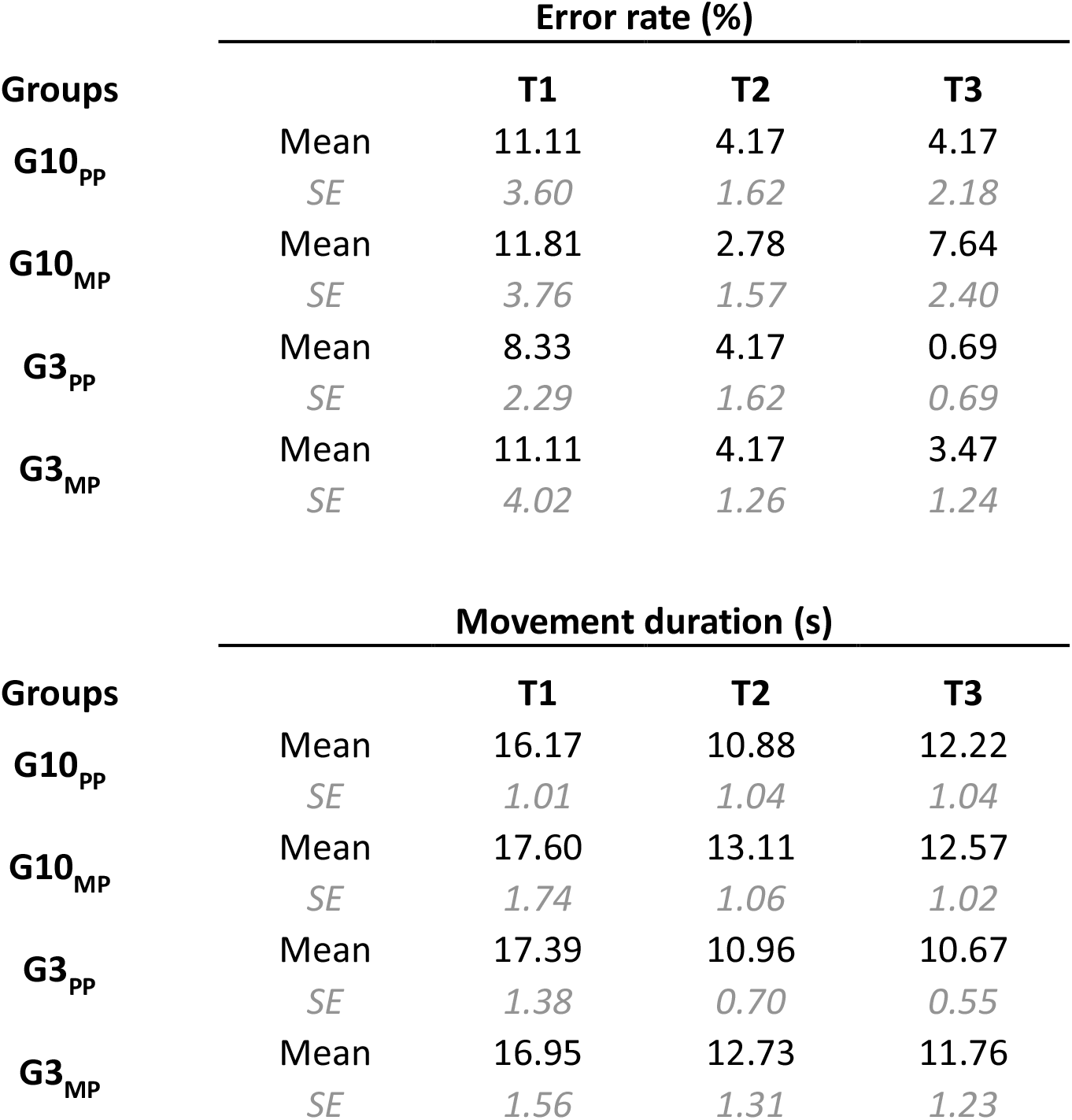
Average value (+SE) of Error rate (%) and Movement duration (s) for the four groups and three sessions.

**Figure 2.**
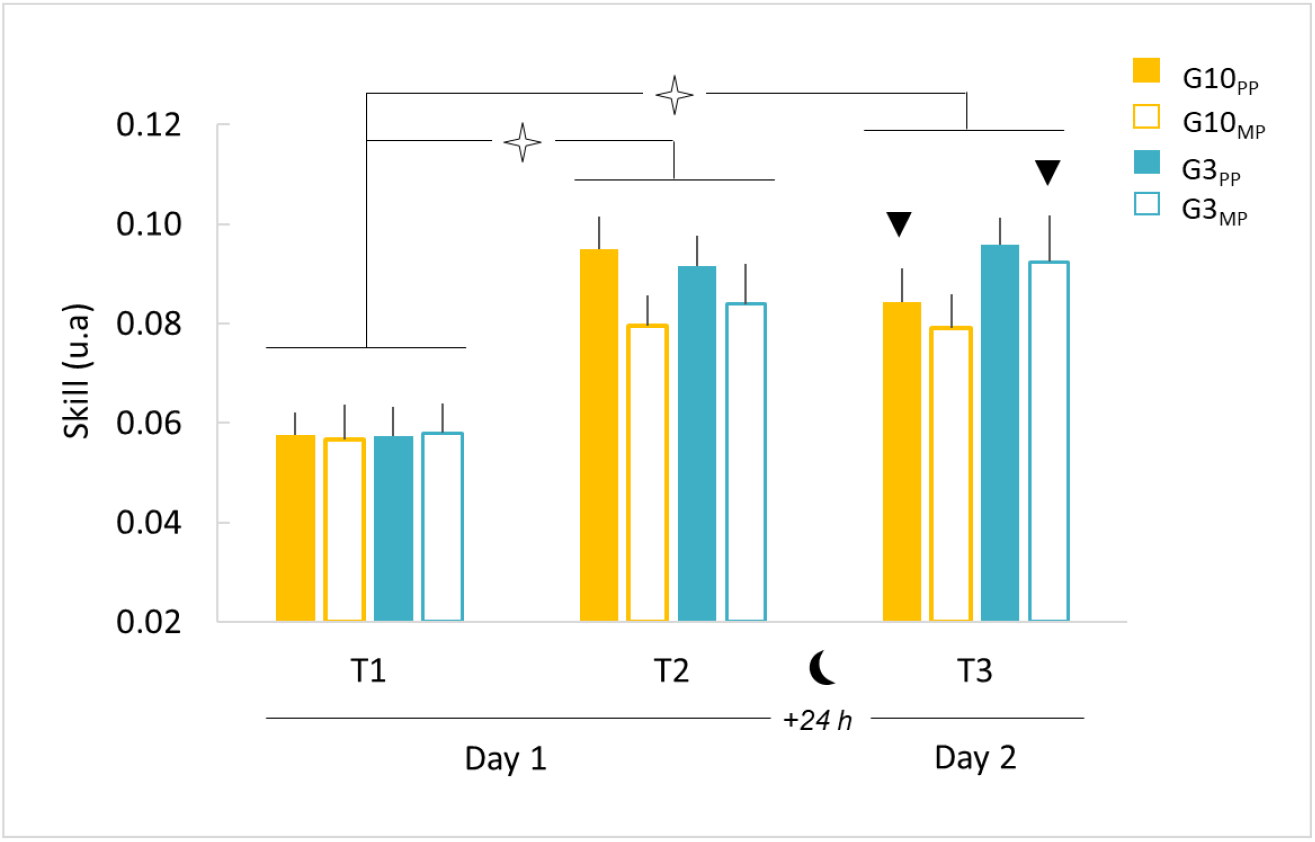
Average values (+SE) of skill performance for the four groups and the three sessions. Open star indicates significant differences between T1 and T2 and between T1 and T3 for all groups. Black triangle indicates significant difference between T2 and T3.

After training, all groups significantly enhanced their skill performance (T1 vs T2; in all, p < 0.001), which was characterized by a diminution of movement duration and error rate (Table 1). One day after training (T2 vs T3), we observed further improvement in skill performance for the G3_MP_ (p = 0.04), a stabilization for the G3_PP_ (p = 0.28) and the G10_MP_ (p = 0.92), and a deterioration for the G10_PP_ (p = 0.01). In detail, the G3_MP_ improved speed and accuracy, the G3_PP_ stabilized speed and improved accuracy, the G10_MP_ improved speed and deteriorated accuracy, while the G10_PP_ deteriorated speed and stabilized accuracy (Table 1). Importantly, despite groups differences in skill improvement after training (T2 vs T3), all groups acquired better skill performance (i.e., consolidation) one day later (T3) compared to their initial performance (T1 vs T3, in all, p < 0.001); see also Table 1 for error rate and movement duration.

To explain in more detail the acquisition and consolidation processes according to the time-of-day and training, we focus on gains between sessions, illustrated in Figure 3 (T1_T2 for acquisition, T2_T3 for consolidation, and T1_T3 for total gain).

**Figure 3.**
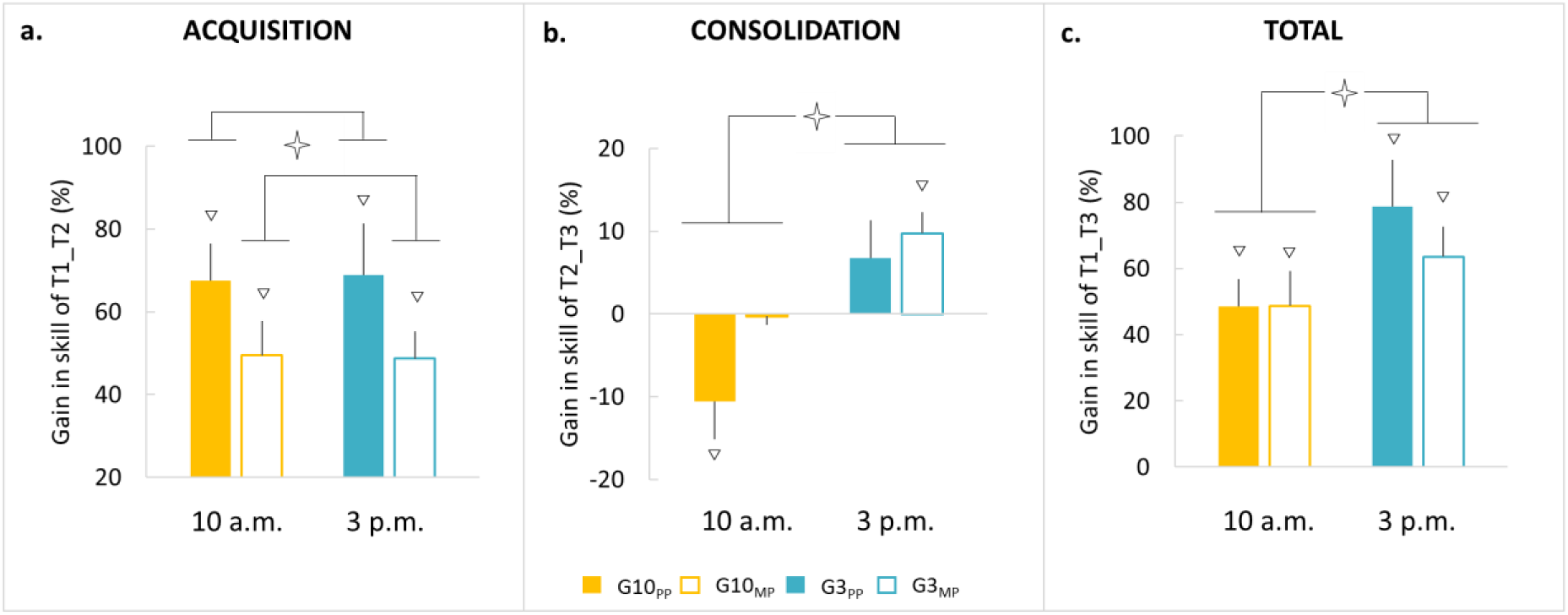
Average values (+SE) of gains in skill performance (%) for the four groups. (a) Acquisition gains (T1_T2). (b) Consolidation gains (T2_T3). (c) Total gains (T1_T3). Open stars indicate significant differences between Practice or Time-of-day. White triangles indicate significant differences from the value *zero*.

### Gain in the acquisition process

On day 1 (Figure 3a), the comparison of T1_T2 gain with the reference value zero (0) showed a significant improvement in skill performance for all groups (in all, t > 5.46; p < 0.001). ANOVA revealed, however, a significant main effect of *practice* (F_1,44_ = 4.10; p < 0.05; η^2^ = 0.08), without *time-of-day* (F_1,44_ = 0.00; p < 0.98) or *interaction* (F_1,44_ = 0.01; p < 0.91) effects, suggesting a better gain following PP than MP, as classically observed in the literature ^3^. Interestingly, the absence of an effect of time-of-day suggests that acquisition processes of mental and physical practices are independent of the time-of-day.

### Gain in the consolidation process

On day 2 (Figure 3b), the comparison of the consolidation gains (T2_T3) with the reference value *zero* (0) showed a deterioration of skill performance for the G10_PP_ (t = -2.27, p = 0.04), a stabilization for the G10_MP_ (t = -0.06, p = 0.95) and the G3_PP_ (t = 1.15, p = 0.16), and an enhancement (i.e., offline learning process) for the G3_MP_ (t = 3.74, p = 0.003). ANOVA showed a significant main effect of *time-of-day* (F_1,44_ = 10.48; p < 0.002; η^2^ = 0.19), without *practice* (F_1,44_ = 2.46; p = 0.12) or *interaction* (F_1,44_ = 0.74; p = 0.39) effects, indicating that the consolidation processes after mental or physical practices was better the afternoon than the morning.

### Total gain

The comparison of the total gains (T1_T3, Figure 3c) with the reference value *zero* (0) showed a significant improvement in skill performance for all groups (in all, t > 4.61, p < 0.001). ANOVA revealed a significant main effect for the *time-of-day* (F_1,44_ = 4.46; p < 0.04; η^2^ = 0.09), without *practice* (F_1,44_ = 0.48; p = 0.49) or *interaction* (F_1,44_ = 0.52; p = 0.47) effect. These results indicated that while all groups acquired better skill performance one day after the training compared to their initial level, the performance was better in the afternoon than the morning, regardless of the type of practice.

## Discussion

We examined the influence of the time-of-day on skill acquisition and consolidation following physical (PP) or mental (MP) practice of a finger-tapping task. Our findings showed a substantial improvement in motor skill after the two types of training (PP and MP) whatever the time of the day (10 a.m. and 3 p.m.); there was, however, better acquisition for the PP compared to MP. Interestingly, we found better consolidation for both PP and MP when the training sessions were scheduled in the afternoon (3 p.m.) compared to the morning (10 a.m.).

### Time-of-day influence on acquisition and consolidation processes

While several studies have reported an influence of the time-of-day on motor performance, such as muscular force ^17^, speed ^28^, or fine motor skills ^18,19^, we did not find such an effect on skill acquisition immediately after PP or MP. Indeed, we found an increase in skill performance following a single session of PP and MP, whatever the time-of-day. In line with our results, Sale et al. have highlighted that the improvement in motor performance following PP is neither influenced by the time-of-day nor by diurnal changes in circulating cortisol levels ^32^. Although not designed to this aim, two previous studies indirectly attained similar conclusions, namely comparable gains in motor performance following PP ^33^ and MP ^34^, whatever the time-of-day of the practice. Overall, neither PP nor MP seems to beneficiate from a particular time during the day to improve skill performance.

Interestingly, we did find a time-of-day effect on skill consolidation. Precisely, one day after the training session, skill performance was further improved when PP or MP took place in the afternoon (3 p.m.) compared to the morning (10 a.m.). Circadian modulations of physiological mechanisms could explain this novel finding. Motor memory formation, following both PP or MP, is associated with neural adaptations within the motor cortex ^35–38^. Intriguingly, Sale et al. suggested that neural plasticity is modulated across the day, due to cortisol hormonal circadian fluctuation ^39^. Indeed, the cortisol concentration, higher in the morning than in the afternoon, was negatively correlated with neural plasticity. Although the influence of the cortisol level on motor consolidation must directly be evaluated, we could speculate that a high level of cortisol during the training would have detrimental effects on the consolidation process. A complementary hypothesis, needing certainly further exploration, implicates the hippocampus. Animal studies have reported that this area is under circadian influence ^40,41^, while the degree of its activation, associated with that of the striatum during PP, seems to predict the performance gain after a night of sleep ^42,43^.

Behaviorally, a possible explanation concerning our findings could be the daily modulation of the sensorimotor predictions of the internal forward models. It has been proposed that during both PP and MP, sensorimotor prediction improves the controller, and thus the motor command ^3,5,9,44,45^. Gueugneau et al. showed a variation of internal predictions across the day, being more accurate in the afternoon than in the morning, which could explain why motor consolidation is better after a practice session in the afternoon ^26,27^. We have also recently demonstrated, using an fMRI experiment, that motor performance is continuously updated daily with a predominant role of the frontoparietal cortex and cerebellum ^29^, which are both involved in the prediction process ^46,47^.

### The differential effects of physical and mental practices on acquisition and consolidation processes

Following previous findings ^3,5,31^, our results showed better acquisition after PP than MP, without time-of-day effects. This difference in acquisition level may be explained by the concept of internal forward models. Evidence supports the hypothesis that internal forward models predict the sensory consequences (e.g., position and velocity) of an upcoming movement, based on the copy of the motor command and the initial state of the apparatus. This prediction is compared with the sensory information from the periphery during the movement. Any discrepancy in this comparison will drive the internal forward model to provide better predictions ^44^ and, in turn, to improve the controller and thus the motor output. A recent study showed that forward models are triggered to predict the sensory consequence of imagined movements ^45^. These internal predictions could improve the motor command in the absence of movement-related sensory feedback ^3,48^. The sensorimotor prediction during imagined movement is, however, more variable ^49^, because it is not updated by sensory feedbacks like physical movement ^50^, which could explain the smaller effectiveness of MP compared to PP in motor performance improvement.

Most interestingly, albeit this difference in the acquisition (i.e., immediately after training), PP and MP obtained similar skill performances one day after the training, with a better total gain in the afternoon than in the morning (see Fig 3c). This adjustment of skill performance between PP and MP could be attributed to a different consolidation process between PP and MP (see Fig 3b). In fact, in the morning, we observed a deterioration of skill performance for the PP versus a stabilization for the MP, suggesting a more efficient consolidation after MP. This forgetting may reflect a fragile memory, more susceptible to interferences, after the acquisition at 10 a.m. for the PP only ^51,52^, while the stabilization of MP reflects a more robust memory ^53^.

Likewise, in the afternoon, MP showed also a more efficient consolidation process, highlighted by an enhancement of performance compared to stabilization for PP. This result corroborates our recent finding for a pointing task ^3^, which showed an enhancement of skill performance 6 hours after MP but not after PP. We explained this result by a slow motor learning process for MP, due to the availability of internal predictions only to drive the controller. Indeed, motor learning through MP may need passage-of-time to be consolidated, while PP may lead to a rapid acquisition with complete consolidation. Thus, our results expand and generalize those of the study of Ruffino et al. ^3^, which suggested that PP and MP involve different acquisition and consolidation processes, leading, however, to similar skill performance one day after the training.

## Conclusion

In conclusion, the present study provides the first evidence of the influence of the time-of-day on the consolidation process following PP or MP. Even if further investigations are required to determine the physiological and/or behavioral bases of these modulations, the findings of the current study have important methodological and practical implications. From a methodological point of view, our data underline the importance to consider the time-of-day when planning experiments investigating motor learning or motor performance. Regarding practical applications, we recommend, when this is possible, to schedule rehabilitation or sports training programs in the afternoon, whatever the type of practice (physical or mental).

## Methods

### Participants

Forty-eight healthy adults participated in the current study after giving their informed consent. All were right-handed (mean score 0.79 ± 0.22), as measured by the Edinburgh handedness questionnaire ^54^, and free from neurological or physical disorders. Participants were randomly assigned into four groups: two PP groups, one trained in the morning (G10_PP_, n = 12, 8 females, mean age: 25 ± 6 years old) and the other trained in the afternoon (G3_PP_, n = 12, 7 females, mean age: 24 ± 6 years old), and two MP groups, one trained in the morning (G10_MP_, n = 12, 3 females, mean age: 25 ± 4 years old) and the other trained in the afternoon (G3_MP_, n = 12, 6 females, mean age: 25 ± 2 years old). Due to the nature of the motor task (finger tapping) used in the present study, we did not include musicians and professional typists. The regional ethic committee (CPP EST) approved the experimental design according to the standards set by the Declaration of Helsinki.

All participants were requested to be drug- and alcohol-free, to not change their habitual daily activities (e.g., cooking, computer use, handiwork), and to not make intensive physical activity during the 24 hours preceding the experiment. They were all synchronized with a normal diurnal activity (8 a.m. ± 1 hour to 12 a.m. ± 1 hour alternating with the night).

We examined the chronotype of each participant using the Morningness-Eveningness Questionnaire ^55^. In this test, scores range from 16 to 86 and are divided in five categories: evening type (score 16 to 30, n = 1), moderate evening type (score 31 to 41, n = 3), intermediate type (score 42 to 58, n = 26), moderate morning type (score 59 to 69, n = 4) and morning type (score 70 to 86, n = 1). One-way ANOVA did not show significant differences between groups (F_3,44_ = 1.81 p = 0.16; mean scores: G10_PP_= 51 ± 11, G10_MP_= 56 ± 12, G3_PP_= 50 ± 8, G3_MP_= 47 ± 9).

We also verified the sleep quality of each participant with the Pittsburgh Sleep Quality Index ^56^. The general score in this questionnaire ranges from 0 (no particular difficulties to sleep) to 21 (major difficulties to sleep). One-way ANOVA indicated very good sleep quality, which was similar between groups (F_3,44_ = 0.47 p = 0.70; mean scores: G10_PP_ = 5 ± 1, G10_MP_= 5 ± 1, G3_PP_= 4 ± 1, G3_MP_= 5 ± 1).

Motor imagery ability for the MP groups was assessed by the French version of the Movement Imagery Questionnaire “MIQr” ^57^. The MIQr is an 8-item self-report questionnaire, in which the participants rate the vividness of their mental images using two 7-point scales, one associated to visual and the other to kinesthetic imagery. The score ‘7’ indicates easy to feel/visualize, whereas the score ‘1’ corresponds to difficult to feel/visualize (maximum score: 56; minimum score: 8). There were no significant differences between the two MP groups (two-tailed t-tests for independent groups; t = 1.11 p = 0.29; mean scores: G10_MP_= 44 ± 7, G3_MP_= 45 ± 5), indicating good imagery ability for each group.

### Experimental device and procedure

We employed a computerized version of the sequential finger-tapping task ^58^, commonly used in laboratory experiments, allowing us to observe online and offline changes in motor performance following motor imagery training ^34^. Participants were comfortably seated on a chair in front of a keyboard. They were requested to tap, as accurately and as fast as possible, with their left hand the following sequence: 1-3-2-4-1-0 (Fig. 1a). Each key was affected to a specific finger: 0 (thumb), 1 (index), 2 (middle), 3 (ring), and 4 (little). One trial was composed of six sequences. Precisely, at the beginning of each trial, participants pressed the key ‘0’ with their thumb to start the chronometer and they accomplished the 6 sequences continuously. Pressing the key ‘0’ at the end of the 6th sequence stopped the chronometer and ended the trial. To familiarize themselves with the protocol, participants accomplished two trials at a natural speed. The vision of the non-dominant hand was hidden through a box during the whole protocol. The sequence’s order, however, was displayed on the box and thus visible to the participants during the whole experiment.

The experiments were scheduled on two consecutive days (Day 1 and Day 2) and at different times within each day (Figure 1b). On Day 1, participants were physically (G10_PP_) or mentally (G10_MP_) trained at 10 a.m. or 3 p.m. (G3_PP_ and G3_MP_, respectively). All participants carried out 48 trials (12 blocks of 4 trials, with 5-s rest between trials and 30-s rest between blocks to avoid mental fatigue ^59^). To evaluate the improvement in skill performance (i.e., the acquisition process) following the two training methods, all groups physically accomplished the first two trials (1 and 2, pre-test, T1) and the last two trials (47 and 48, post-test, T2). The remaining trials (3-46, n = 44) constituted the training trials for the physical (G10_PP_ and G3_PP_) or the mental (G10_MP_ and G3_MP_) groups. To ensure that all participants of G10_MP_ and G3_MP_ would correctly complete mental training, we provided the following instructions: ‘try to imagine yourself performing the motor task, by seeing and feeling your arm moving as if you were actually moving it’. To test the consolidation process, the participants of each group performed two trials twenty-four hours after the end of the training (T3). Note that no instructions concerning the motor performance (i.e., speed or typing errors) were provided to the participants.

### Data recording and analysis

For T1, T2, and T3, a program Visual Basic for Applications (Microsoft, Excel) recorded the accuracy and movement duration in the pre-test and post-test ^3^. The accuracy (error rate) was defined as the number of false sequences throughout one trial (0 = no error during the trial; 6 = maximum number of errors). If the participants made one or more mistakes in one of the sequences, we counted this sequence as false (see Fig. 1a). The error rate was defined as the percentage of the number of errors during a trial:

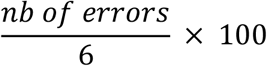

Movement duration was defined as the time interval between the start of the trial (when the participants pressed the first key ‘0’) and the end of the trial (when the participants pressed the key ‘0’ at the end of the 6th sequence).

These two parameters (movement duration and error rate) are related by the speed-accuracy tradeoff function ^60^. Ascertaining that motor skill (i.e., the training-related change in the speed-accuracy trade-off function) has been improved, duration and accuracy should not change in opposite directions. For that reason, we compute a composite ratio of duration and accuracy to describe motor skill as follows:

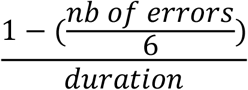

Note that skill increases when the ratio increases.

Gains for the acquisition and consolidation were calculated as follows:

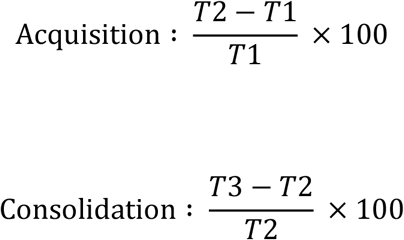

The total gain was calculated as follows:

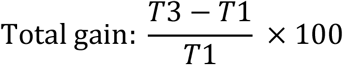

Finally, to verify that participants did not activate their muscles during MP electromyographic (EMG) activity of the first dorsal interosseous (FDI) of the left hand was recording during each imagined movement and compared to EMG activity at rest (10 seconds recording before training). We used a pair of bipolar silver chloride circular (recording diameter of 10 mm) surface electrodes positioned lengthwise over the middle of the muscles belly with an interelectrode (center to center) distance of 20 mm. The reference electrode was placed on the medial elbow epicondyle. After shaving and dry-cleaning the skin with alcohol, the impedance was below 5 kΩ. EMG signals were amplified (gain 1000), filtered (with a bandwidth frequency ranging from 10 Hz to 1 kHz), and converted for digital recording and storage with PowerLab 26T and LabChart 7 (AD Instruments). We analyzed the EMG patterns of the muscle by computing their activation level (RMS, root mean square) using the following formula:

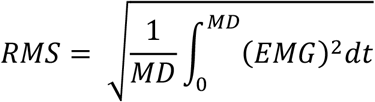

The statistical analysis did not show any significant difference between the EMG recording during motor imagery and the EMG recording at rest (in all, p > 0.05).

### Statistical analysis

Statistical analyses were completed using STATISTICA (8.0 version: Stat-Soft, Tulsa, OK). The normality of the data distributions and sphericity was verified using Shapiro-Wilk (p > 0.05) and Mauchly’s test (p < 0.05), respectively.

As a first step, analyses were performed to control for potential methodological biases. We compared the chronotype and the quality of sleep between groups (G10_PP_, G10_MP_, G3_PP,_ and G3_MP_) with one-way ANOVA. Also, we compared the motor imagery capacities between the two MP groups (G10_MP_ vs G3_MP_) with a two-tailed independent samples *t-test*.

Then, we applied repeated measures (rm) ANOVA with *group* as between-subject factor (G10_PP_, G10_MP_, G3_PP,_ and G3_MP_) and *session* as the within-subject factor (T1, T2, and T3) for motor skill. To further assess the influence of the type of practice and the time-of-day on acquisition and consolidation processes, we conducted two factorial ANOVA on T1_T2 and T2_T3 skill gains with categorical factors “Practice” (PP vs. MP) and “Time-of-day” (Morning vs. Afternoon). Finally, to analyze the total gain in skill performance (gain between T1 and T3), we performed a third factorial ANOVA with the same factors. All post-hoc analyses were performed by applying Fischer’s tests. A supplementary *t-test* analysis permitted us to compare each gain (acquisition, consolidation, and total) with the reference value zero (0) for each group.

## Data availability

The datasets generated during and analyzed during the current study are available from the corresponding author on reasonable request.

## Author contributions

CT, CP, and CR designed the experiment; CT, FB, and CR recorded the data; CT and PMH analyzed the data; CT developed figures; CT, PMH, CP, and CR wrote the manuscript; CP, PHM, MB, FL, and CR provided feedback on the manuscript; all co-authors read and approved the submitted version.

## References

1. Doyon, J. & Benali, H. Reorganization and plasticity in the adult brain during learning of motor skills. Curr. Opin. Neurobiol. 15, 161–167 (2005).

2. Walker, M. P., Brakefield, T., Morgan, A., Hobson, J. A. & Stickgold, R. Practice with sleep makes perfect: Sleep-dependent motor skill learning. Neuron 35, 205–211 (2002).

3. Ruffino, C. et al. Acquisition and consolidation processes following motor imagery practice. Sci. Rep. 11, 2295 (2021).

4. Tunovic, S., Press, D. Z. & Robertson, E. M. A Physiological Signal That Prevents Motor Skill Improvements during Consolidation. J. Neurosci. 34, 5302–5310 (2014).

5. Gentili, R., Han, C. E., Schweighofer, N. & Papaxanthis, C. Motor Learning Without Doing: Trial-by-Trial Improvement in Motor Performance During Mental Training. J. Neurophysiol. 104, 774–783 (2010).

6. Ruffino, C., Papaxanthis, C. & Lebon, F. Neural plasticity during motor learning with motor imagery practice: Review and perspectives. Neuroscience 341, 61–78 (2017).

7. Lebon, F., Collet, C. & Guillot, A. Benefits of Motor Imagery Training on Muscle Strength. J. Strength Cond. Res. 24, 1680–1687 (2010).

8. Grosprêtre, S., Jacquet, T., Lebon, F., Papaxanthis, C. & Martin, A. Neural mechanisms of strength increase after one-week motor imagery training. Eur. J. Sport Sci. 18, 209–218 (2018).

9. Gentili, R., Papaxanthis, C. & Pozzo, T. Improvement and generalization of arm motor performance through motor imagery practice. Neuroscience 137, 761–772 (2006).

10. Page, S. J., Szaflarski, J. P., Eliassen, J. C., Pan, H. & Cramer, S. C. Cortical Plasticity Following Motor Skill Learning During Mental Practice in Stroke. Neurorehabil. Neural Repair 23, 382–388 (2009).

11. Shoenfelt, E. L. & Griffith, A. U. Evaluation of a Mental Skills Program for Serving for an Intercollegiate Volleyball Team. Percept. Mot. Skills 107, 293–306 (2008).

12. Guillot, A. & Collet, C. Duration of Mentally Simulated Movement: A Review. J. Mot. Behav. 37, 10–20 (2005).

13. Hétu, S. et al. The neural network of motor imagery: An ALE meta-analysis. Neuroscience and Biobehavioral Reviews vol. 37 930–949 (2013).

14. Grosprêtre, S., Lebon, F., Papaxanthis, C. & Martin, A. New evidence of corticospinal network modulation induced by motor imagery. J. Neurophysiol. 115, 1279–1288 (2016).

15. Grosprêtre, S., Lebon, F., Papaxanthis, C. & Martin, A. Spinal plasticity with motor imagery practice. J. Physiol. 597, 921–934 (2019).

16. Di Rienzo, F. et al. Online and Offline Performance Gains Following Motor Imagery Practice: A Comprehensive Review of Behavioral and Neuroimaging Studies. Front. Hum. Neurosci. 10, 315 (2016).

17. Guette, M., Gondin, J. & Martin, A. Time-of-Day Effect on the Torque and Neuromuscular Properties of Dominant and Non-Dominant Quadriceps Femoris. Chronobiol. Int. 22, 541–558 (2005).

18. Jasper, I., Häubler, A., Marquardt, C. & Hermsdörfer, J. Circadian rhythm in handwriting. J. Sleep Res. 18, 264–271 (2009).

19. Edwards, B., Waterhouse, J. & Reilly, T. The Effects of Circadian Rhythmicity and Time-Awake on a Simple Motor Task. Chronobiol. Int. 24, 1109–1124 (2007).

20. Reilly, T. & Down, A. Investigation of circadian rhythms in anaerobic power and capacity of the legs. J. Sports Med. Phys. Fitness 32, 343–347 (1992).

21. Atkinson, G. & Speirs, L. Diurnal Variation in Tennis Service. Percept. Mot. Skills 86, 1335–1338 (1998).

22. Edwards, B. J., Lindsay, K. & Waterhouse, J. Effect of time of day on the accuracy and consistency of the badminton serve. Ergonomics 48, 1488–1498 (2005).

23. Kline, C. E. et al. Circadian variation in swim performance. J. Appl. Physiol. 102, 641–649 (2007).

24. Atkinson, G. & Reilly, T. Circadian Variation in Sports Performance. Sport. Med. 21, 292–312 (1996).

25. Thun, E., Bjorvatn, B., Flo, E., Harris, A. & Pallesen, S. Sleep, circadian rhythms, and athletic performance. Sleep Med. Rev. 23, 1–9 (2015).

26. Gueugneau, N. & Papaxanthis, C. Time-of-day effects on the internal simulation of motor actions: psychophysical evidence from pointing movements with the dominant and non-dominant arm. Chronobiol. Int. 27, 620–639 (2010).

27. Gueugneau, N., Schweighofer, N. & Papaxanthis, C. Daily update of motor predictions by physical activity. Sci. Rep. 5, 17933 (2015).

28. Gueugneau, N., Pozzo, T., Darlot, C. & Papaxanthis, C. Daily modulation of the speed– accuracy trade-off. Neuroscience 356, 142–150 (2017).

29. Bonzano, L., Roccatagliata, L., Ruggeri, P., Papaxanthis, C. & Bove, M. Frontoparietal cortex and cerebellum contribution to the update of actual and mental motor performance during the day. Sci. Rep. 6, 30126 (2016).

30. Allami, N., Paulignan, Y., Brovelli, A. & Boussaoud, D. Visuo-motor learning with combination of different rates of motor imagery and physical practice. Exp. Brain Res. 184, 105–113 (2008).

31. Gentili, R. J. & Papaxanthis, C. Laterality effects in motor learning by mental practice in right-handers. Neuroscience 297, 231–242 (2015).

32. Sale, M. V., Ridding, M. C. & Nordstrom, M. A. Time of Day Does Not Modulate Improvements in Motor Performance following a Repetitive Ballistic Motor Training Task. Neural Plast. 2013, 1–9 (2013).

33. Walker, M. P. Sleep and the Time Course of Motor Skill Learning. Learn. Mem. 10, 275–284 (2003).

34. Debarnot, U., Creveaux, T., Collet, C., Doyon, J. & Guillot, A. Sleep Contribution to Motor Memory Consolidation: A Motor Imagery Study. Sleep 32, 1559–1565 (2009).

35. Pascual-Leone, A. et al. Modulation of muscle responses evoked by transcranial magnetic stimulation during the acquisition of new fine motor skills. J. Neurophysiol. 74, 1037–1045 (1995).

36. Rosenkranz, K., Kacar, A. & Rothwell, J. C. Differential Modulation of Motor Cortical Plasticity and Excitability in Early and Late Phases of Human Motor Learning. J. Neurosci. 27, 12058–12066 (2007).

37. Avanzino, L. et al. Motor cortical plasticity induced by motor learning through mental practice. Front. Behav. Neurosci. 9, 105 (2015).

38. Ruffino, C., Gaveau, J., Papaxanthis, C. & Lebon, F. An acute session of motor imagery training induces use-dependent plasticity. Sci. Rep. 9, 20002 (2019).

39. Sale, M. V., Ridding, M. C. & Nordstrom, M. A. Cortisol Inhibits Neuroplasticity Induction in Human Motor Cortex. J. Neurosci. 28, 8285–8293 (2008).

40. Devan, B. D. et al. Circadian Phase-Shifted Rats Show Normal Acquisition but Impaired Long-Term Retention of Place Information in the Water Task. Neurobiol. Learn. Mem. 75, 51–62 (2001).

41. Smarr, B. L., Jennings, K. J., Driscoll, J. R. & Kriegsfeld, L. J. A time to remember: The role of circadian clocks in learning and memory. Behav. Neurosci. 128, 283–303 (2014).

42. Albouy, G. et al. Maintaining vs. enhancing motor sequence memories: Respective roles of striatal and hippocampal systems. Neuroimage 108, 423–434 (2015).

43. King, B. R., Gann, M. A., Mantini, D., Doyon, J. & Albouy, G. Persistence of Hippocampal Multivoxel Patterns during Awake Rest after Motor Sequence Learning. bioRxiv 1–21 (2021) doi:10.1101/2021.06.29.450290.

44. Wolpert, D. M., Ghahramani, Z. & Flanagan, J. R. Perspectives and problems in motor learning. Trends Cogn. Sci. 5, 487–494 (2001).

45. Kilteni, K., Andersson, B. J., Houborg, C. & Ehrsson, H. H. Motor imagery involves predicting the sensory consequences of the imagined movement. Nat. Commun. 9, 1617 (2018).

46. Wolpert, D. M., Miall, R. C. & Kawato, M. Internal models in the cerebellum. Trends Cogn. Sci. 2, 338–347 (1998).

47. Blakemore, S.-J. & Sirigu, A. Action prediction in the cerebellum and in the parietal lobe. Exp. Brain Res. 153, 239–245 (2003).

48. Ingram, T. G. J., Solomon, J. P., Westwood, D. A. & Boe, S. G. Movement related sensory feedback is not necessary for learning to execute a motor skill. Behav. Brain Res. 359, 135–142 (2019).

49. Courtine, G., Papaxanthis, C., Gentili, R. & Pozzo, T. Gait-dependent motor memory facilitation in covert movement execution. Cogn. Brain Res. 22, 67–75 (2004).

50. Demougeot, L. & Papaxanthis, C. Muscle Fatigue Affects Mental Simulation of Action. J. Neurosci. 31, 10712–10720 (2011).

51. Korman, M. et al. Daytime sleep condenses the time course of motor memory consolidation. Nat. Neurosci. 10, 1206–1213 (2007).

52. Robertson, E. M. New Insights in Human Memory Interference and Consolidation. Curr. Biol. 22, R66–R71 (2012).

53. Debarnot, U., Maley, L., Rossi, D. D. & Guillot, A. Motor interference does not impair the memory consolidation of imagined movements. Brain Cogn. 74, 52–57 (2010).

54. Oldfield, R. C. The assessment and analysis of handedness: The Edinburgh inventory. Neuropsychologia 9, 97–113 (1971).

55. Horne, J. A. & Ostberg, O. A self assessment questionnaire to determine Morningness Eveningness in human circadian rhythms. Int. J. Chronobiol. 4, 97–110 (1976).

56. Buysse, D. J., Reynolds, C. F., Monk, T. H., Berman, S. R. & Kupfer, D. J. The Pittsburgh sleep quality index: A new instrument for psychiatric practice and research. Psychiatry Res. 28, 193–213 (1989).

57. Lorant, J. & Nicolas, A. Validation de la traduction française du Movement Imagery Questionnaire-Revised (MIQ-R). Sci. Mot. 53, 57–68 (2004).

58. Kami, A. et al. Functional MRI evidence for adult motor cortex plasticity during motor skill learning. Nature 377, 155–158 (1995).

59. Rozand, V., Lebon, F., Stapley, P. J., Papaxanthis, C. & Lepers, R. A prolonged motor imagery session alter imagined and actual movement durations: Potential implications for neurorehabilitation. Behav. Brain Res. 297, 67–75 (2016).

60. Shmuelof, L., Krakauer, J. W. & Mazzoni, P. How is a motor skill learnedã Change and invariance at the levels of task success and trajectory control. J. Neurophysiol. 108, 578–594 (2012).

